# The Renin–Angiotensin System Modulates SARS-CoV-2 Entry via ACE2 Receptor

**DOI:** 10.1101/2025.06.25.661409

**Authors:** Sophia Gagliardi, Tristan Hotchkin, Hasset Tibebe, Grace Hillmer, Dacia Marquez, Coco Izumi, Jason Chang, Alexander Diggs, Jiro Ezaki, Yuichiro J. Suzuki, Taisuke Izumi

## Abstract

The RAS plays a central role in cardiovascular regulation and has gained prominence in the pathogenesis of COVID-19 due to the critical function of ACE2 as the entry receptor for SARS-CoV-2. Angiotensin IV, but not angiotensin II, has recently been reported to enhance the binding between the viral spike protein and ACE2. To investigate the virological significance of this effect, we developed a single-round infection assay using SARS-CoV-2 viral-like particles expressing the spike protein. Our results demonstrate that while angiotensin II does not affect viral infectivity across concentrations ranging from 40 nM to 400 nM, angiotensin IV enhances viral entry at a low concentration but exhibits dose-dependent inhibition at higher concentrations. These findings highlight the unique dual role of angiotensin IV in modulating SARS-CoV-2 entry. In silico molecular docking simulations indicate that angiotensin IV was predicted to associate with the S1 domain near the receptor-binding domain in the open spike conformation. Given that reported plasma concentrations of angiotensin IV range widely from 17 pM to 81 nM, these levels may be sufficient to promote, rather than inhibit, SARS-CoV-2 infection. This study identifies a novel link between RAS-derived peptides and SARS-CoV-2 infectivity, offering new insights into COVID-19 pathophysiology and informing potential therapeutic strategies.

## 1. Introduction

Severe Acute Respiratory Syndrome Coronavirus 2 (SARS-CoV-2), the causative agent of Coronavirus Disease 2019 (COVID-19), is an enveloped, positive-sense, single-stranded RNA virus belonging to the Betacoronavirus genus [1, 2]. Its genome encodes four major structural proteins, nucleocapsid (N), membrane (M), envelope (E), and spike (S), as well as 16 non-structural proteins involved in viral replication and host interaction [3]. Among these, the spike glycoprotein plays a pivotal role in viral entry by mediating host cell recognition and membrane fusion. This protein comprises two functional subunits: S1, which contains the receptor-binding domain (RBD) responsible for engaging host receptors, and S2, which facilitates viral and host membrane fusion. The primary receptor for SARS-CoV-2 entry is angiotensin-converting enzyme 2 (ACE2), a surface carboxypeptidase expressed in various human tissues [4, 5]. Upon binding of the RBD spike protein to ACE2, the spike protein undergoes proteolytic activation by host proteases such as TMPRSS2, triggering conformational changes that allow the virus to penetrate the host cell [4, 5]. In the absence or low expression of TMPRSS2, SARS-CoV-2 can instead utilize an endocytic pathway, where the spike protein is cleaved by cathepsin L within the acidic environment of endosomes to enable membrane fusion and entry [6]. Despite the central role of ACE2 receptor in viral entry, its expression is variable across tissues, and relatively low in the lungs, prompting ongoing investigation into factors that modulate viral access and entry efficiency [7]. ACE2 is also a critical regulator of the renin-angiotensin system (RAS), a hormonal cascade essential for cardiovascular and renal homeostasis [8, 9]. Within this system, angiotensinogen is cleaved by renin to form angiotensin I, which is subsequently converted to angiotensin II by ACE (Figure 1). Angiotensin II primarily signals through the angiotensin II type 1 receptor (AT1R), eliciting vasoconstriction, pro-inflammatory responses, and fluid retention [10]. However, angiotensin II can be further processed by aminopeptidases to generate downstream metabolites with distinct biological roles.

**Figure 1.**
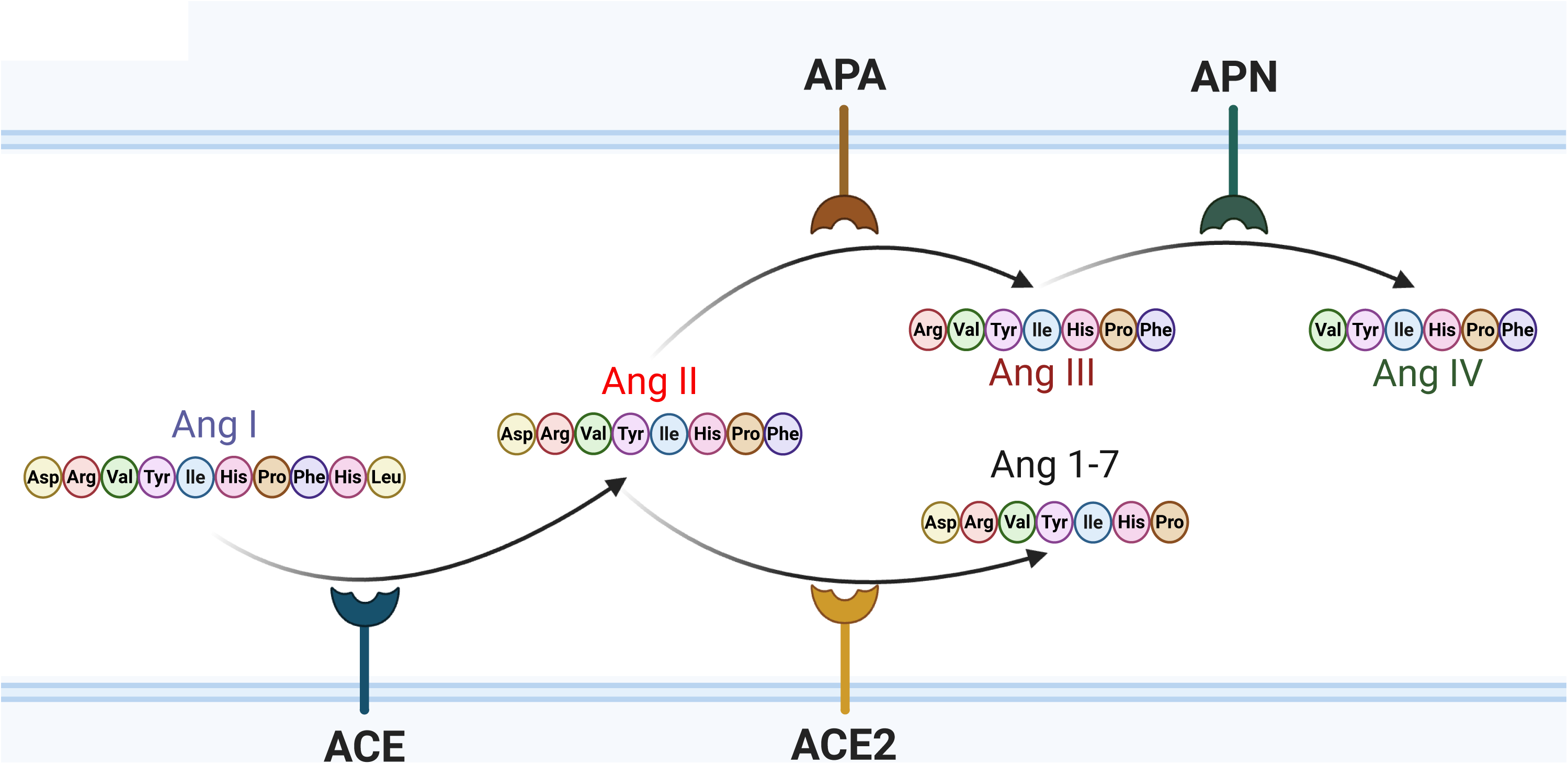
Schematic Representation of Amino Acid Sequences and Metabolic Pathways of Angiotensin Peptides. Angiotensin I (Ang I) is converted to angiotensin II (Ang II) by angiotensin-converting enzyme (ACE). Ang II can be further metabolized by ACE2 to produce angiotensin 1–7 (Ang 1–7), or by aminopeptidase A (APA) to generate angiotensin III (Ang III). Ang III is subsequently cleaved by aminopeptidase N (APN) to produce angiotensin IV (Ang IV).

Aminopeptidase A (APA) cleaves Angiotensin II to produce angiotensin III, which retains affinity for both AT1R and the angiotensin II type 2 receptor (AT2R), but with reduced potency [11]. Angiotensin III is further degraded by aminopeptidase N (APN) into angiotensin IV, a peptide that exerts effects through the AT4 receptor (AT4R), also known as insulin-regulated aminopeptidase (IRAP) [12, 13]. AT4R is implicated in diverse physiological functions, including natriuresis, vasodilation, and cognitive processes such as learning and memory [14, 15]. Given the dual role of ACE2 in RAS regulation and as the SARS-CoV-2 entry receptor, interactions between angiotensin peptides and viral entry mechanisms are of growing interest. Recent studies have suggested a complex interplay between the hormonal milieu of the RAS and SARS-CoV-2 infection. The spike protein leads to ACE2 downregulation via interaction, which contributes to the loss of ACE2-induced protective effects and stimulation of counter-regulatory angiotensin II-induced pathogenesis, promoting vasoconstriction, inflammation, and oxidative stress, thereby exacerbating COVID-19 pathogenesis [16]. Mathematical models have suggested that SARS-CoV-2-induced ACE2 downregulation perturbs RAS homeostasis, and interventions targeting RAS components predicted an improvement of the clinical outcome of COVID-19 for some drugs and a worsening for others [17]. This indicates that SARS-CoV-2 infection also influences the balance of the RAS cascade, which leads to severe COVID-19 diseases. It has been reported that ACE2 expression is higher in females than in males, leading to increased circulating levels of angiotensin (1–7) in females [18, 19], while angiotensin II tends to be higher in males [20–22]. When comparing COVID-19 outcomes between males and females, a clear trend emerges that males are hospitalized more frequently and are more likely to require intensive interventions such as intubation and vasopressor support [23, 24]. Furthermore, hospitalized male patients face a higher risk of thrombosis and mortality compared to their female counterparts [25]. The underlying causes of these disparities remain unclear, but they may be attributed to various biological differences between the sexes, and one potential factor may be the differences in circulating angiotensin peptides and their biological activities.

Recent findings suggest that angiotensin peptides may influence not only COVID-19 complications but also viral infectivity. ELISA-based spike protein binding assays have shown that while angiotensin II has no significant effect, the smaller downstream metabolite angiotensin IV enhances the binding affinity of the SARS-CoV-2 spike protein to the ACE2 receptor in vitro [26]. In this study, we investigated the virological implications of angiotensin IV on SARS-CoV-2 infectivity and found that enhanced spike–ACE2 interaction correlates with increased viral entry at lower tested concentrations of angiotensin IV. However, higher concentrations of angiotensin IV dose-dependently inhibited viral entry, indicating that its role in infection is more complex. As spike-mediated downregulation of ACE2 may shift angiotensin II metabolism toward increased production of angiotensin III via APA, and subsequently angiotensin IV, rather than angiotensin (1–7) (Figure 1), this metabolic shift could establish a regulatory feedback loop. Given the reported plasma concentrations of angiotensin IV ranging from approximately 17 pM to 80 nM [27, 28], it is more likely to enhance SARS-CoV-2 infectivity rather than inhibit it, potentially compensating for spike protein–mediated downregulation of ACE2 on target cells. Furthermore, the greater stability and abundance of angiotensin IV in the central nervous system (CNS) suggests that it may predominantly contribute to the neurological complications associated with COVID-19.

## 2. Materials and Methods

### 2.1. Peptides

Angiotensin II was purchased from APExBIO Technology LLC (Houston, TX, USA). Angiotensin IV was purchased from Bachem Americas, Inc. (Torrance, CA, USA). The peptides were dissolved in deionized water at a concentration of 2 mM to prepare stock solutions and were subsequently diluted in phosphate-buffered saline (PBS) (CELLTREAT, Ayer, MA, USA) to the appropriate working concentrations for each experiment.

### 2.2. Plasmid DNAs

The plasmid DNAs, CoV2-N-WT-Hu1, CoV2-M-IRES-E, and Luc-PS9, originally developed by Doudna’s lab at the University of California, Berkeley [29], were obtained from Addgene (Watertown, MA, USA). The plasmid pcDNA3.1-SARS-S2, developed by the Fang Li lab at the University of Minnesota [30], was also obtained from Addgene.

### 2.3. Cell Culture

HEK293T cells were cultured in Dulbecco’s Modified Eagle’s Medium (DMEM; Cytiva, Marlborough, MA, USA) supplemented with 10% fetal bovine serum (FBS; Gibco, Waltham, MA, USA), 1% penicillin–streptomycin–glutamine (Gibco, Waltham, MA, USA), and 1% GlutaMAX (Gibco, Waltham, MA), referred to as D10 medium. Cells were maintained in 100-mm cell culture dishes (CELLTREAT, Ayer, MA, USA) at the manufacturer-recommended seeding density, incubated in the CO_2_ incubator at 37°C in a 5% CO₂ environment. HEK293T-ACE2 stable cells (NR-52511), obtained from BEI Resources (NIAID, NIH), operated by ATCC, were cultured under the same conditions in 35 mm cell culture dishes using D10 medium. For the post-infection angiotensin IV treatment, infected HEK293T cells were washed once with D10 medium and then cultured in fresh D10 medium containing 40 nM angiotensin IV for an additional 24 hours in a CO₂ incubator.

### 2.4. Immunostaining

HEK293T-ACE2 cells were stained with Alexa Fluor 647-conjugated anti-human ACE2 antibody (R&D Systems, Minneapolis, MN, USA) at a concentration of 1 µL per million cells diluted with staining buffer (Invitrogen, Carlsbad, CA, USA), following pre-incubation with 10% Normal Mouse IgG (Invitrogen, Carlsbad, CA, USA) to block Fc-receptor for non-specific binding. Viability was assessed using the Live/Dead Fixable Red Cell Stain Kit for 638 nm excitation (Invitrogen, Carlsbad, CA, USA), and cells were subsequently fixed with 2% formaldehyde (Cell Signaling Technology, Danvers, MA, USA). Immediately after fixation, samples were analyzed on a CytoFLEX flow cytometer (Beckman Coulter, Brea, CA, USA). All staining procedures were performed according to the manufacturers’ instructions.

### 2.5. SARS-CoV-2 viral-like particle (VLP) production

HEK293T cells, seeded at 7 × 10L per 100 mm cell culture dish pre-coated with 0.01% poly-L-lysine (Sigma, St. Louis, MO, USA), were transfected with four SARS-CoV-2 structure proteins encoding plasmid DNAs, CoV2-N-WT-Hu1, CoV2-M-IRES-E, pcDNA3.1-SARS-S2, and Luc-PS9, at a 1:1:1:1 mass ratio using polyethyleneimine (PEI) transfection reagent (Polysciences, Warrington, PA, USA). Three hours after adding the PEI–DNA mixture, the culture medium was replaced with fresh D10 medium. After 48 hours, the virus-containing supernatant was harvested and filtered through a 0.45 µm sterile polyvinylidene difluoride (PVDF) membrane filter. The filtered viral supernatant was concentrated up to 10-fold using polyethylene glycol 8000 (PEG-8000: PROMEGA, Madison, WI, USA) by mixing the supernatant with PEG-8000 at a 3:1 volume ratio. The mixture was gently rotated at 60 rpm for 4 hours and then centrifuged at 1,600 × g for 1 hour at 4 °C. The pellet (SARS2-VLP) was resuspended in fresh D10 medium to achieve a 10-fold concentration and stored at 4 °C until use in infection assays.

### 2.6. Single-Round Infection Assay

HEK293T-ACE2 cells were seeded at 30,000 cells per well in a 96-well flat-bottom plate (CELLTREAT, Ayer, MA, USA) pre-coated with 0.01% poly-L-lysine (Sigma, St. Louis, MO, USA). Each well was then treated with 200 µL of concentrated SARS2-VLP. At 24 hours post-infection, the culture medium containing VLPs was removed, and each well was washed once with 200 µL of D10 medium before replacing with fresh D10. At 48 hours post-infection, luciferase activity was measured according to the manufacturer’s instructions (Promega Corporation, Madison, WI, USA) in a Molecular Devices SpectraMax Microplate Reader (Molecular Devices, San Jose, CA, USA).

### 2.7. In Silico Docking Simulation

Protein–ligand docking simulations were conducted to evaluate interactions between angiotensin peptides and SARS-CoV-2 spike protein structures. The crystal structures used were the RBD of the SARS-CoV-2 spike protein bound to ACE2 (PDB ID: 6M0J) [31] and the trimeric closed and opened forms of the spike glycoprotein (PDB ID: 6VXX and 6VYB, respectively) [32]. Simulations were performed using DynamicBind [33], a deep equivariant generative model for predicting ligand-specific protein–ligand complex structures, accessed via the DynamicBind web server on NeuroSnap. Structural data for angiotensin II (PubChem CID: 172198) and angiotensin IV (PubChem CID: 123814) were provided in SDF file format for input into the simulations. Structural visualizations and images were generated and exported using PyMOL, a user-supported molecular visualization system.

### 2.8. Statistical Analysis

Statistical significance was assessed using the Wilcoxon matched-pairs signed-rank test, comparing each experimental condition to its respective control group.

## 3. Results

### 3.1. Impact of Angiotensin Peptides on SARS-CoV-2 Cell Entry Through ACE2 Receptor

To evaluate the virological relevance of the RAS in SARS-CoV-2 infectivity, we conducted a single-round infection assay using SARS-CoV-2 virus-like particles (SARS2-VLPs) engineered to express a luciferase reporter gene (Figure 2). First, we confirmed that parental HEK293T cells do not express endogenous ACE2, as determined by flow cytometry (Figure 2A). In contrast, HEK293T cells stably expressing ACE2 (HEK293T-ACE2) exhibited robust surface expression, with over 90% of the cell population positively expressing the ACE2 receptor (Figure 2A). Next, we confirmed that our single-round infection assay is ACE2-dependent by using HEK293T cells as target cells. While SARS2-VLPs failed to infect parental HEK293T cells, a significant increase in luciferase activity was observed in HEK293T-ACE2 cells following infection (Figure 2B), indicating that SARS2-VLP target cell entry occurs in an ACE2-dependent manner.

**Figure 2.**
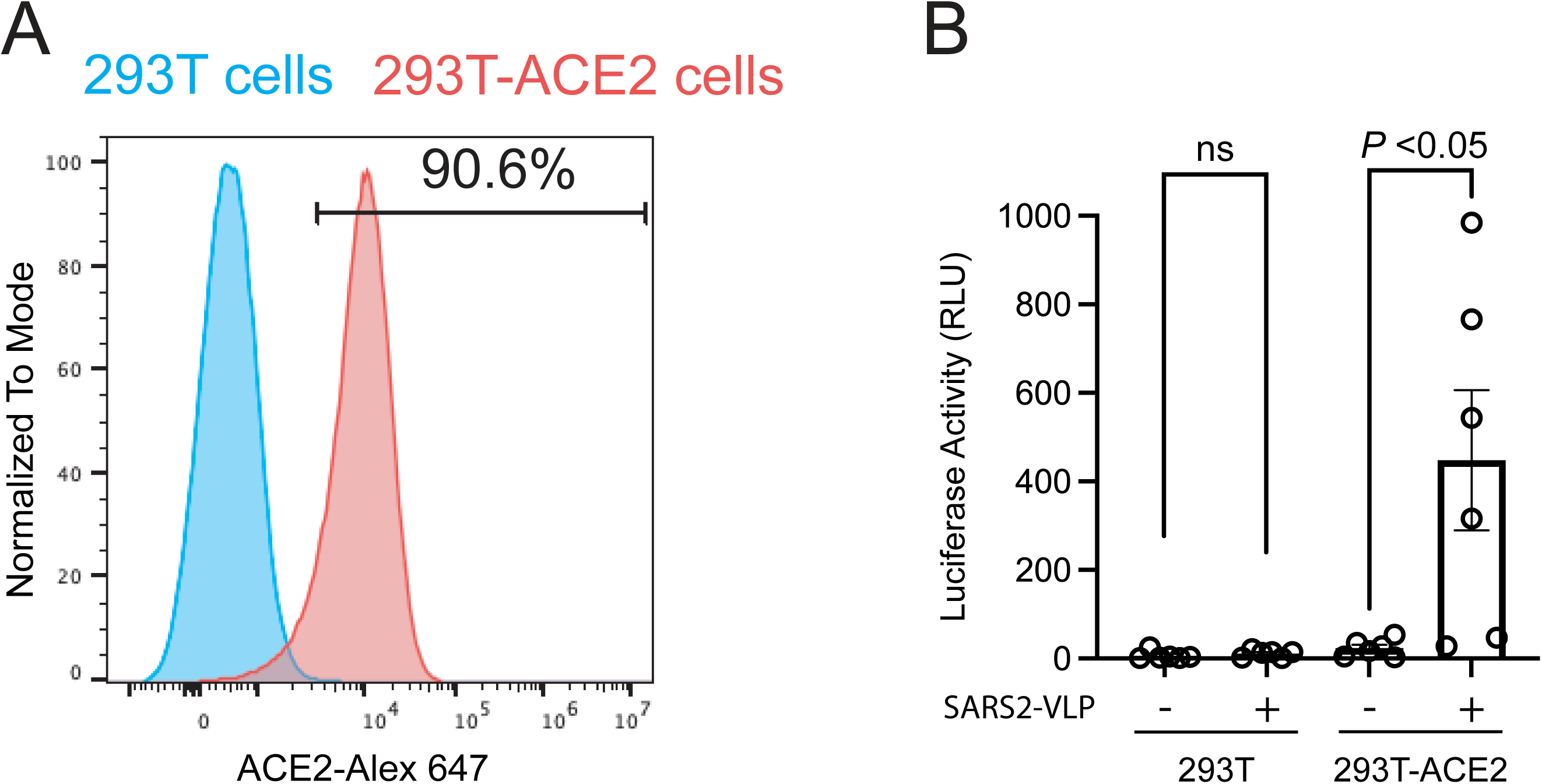
Single-round infection assay with SARS-CoV-2 VLP in 293T-ACE2 cells. **(A)** Flow cytometry analysis confirming ACE2 receptor expression in HEK293T-ACE2 cells used as target cells in the infection assay. HEK293T-ACE2 and parental HEK293T cells were stained with an Alexa Fluor 647-conjugated anti-ACE2 antibody. Parental HEK293T cells showed no detectable ACE2 expression, whereas HEK293T-ACE2 cells exhibited robust surface ACE2 expression. **(B)** Viral infectivity was assessed in HEK293T-ACE2 cells and parental HEK293T cells using a luciferase reporter assay to confirm that SARS-CoV-2 VLP infection is ACE2-dependent. Error bars indicate standard error from six samples and statistical significance was calculated by the Wilcoxon matched-pairs signed-rank test.

Based on the reported ELISA assay that involved pretreatment of the spike protein with angiotensin peptides to assess the enhancement of spike protein interactions with ACE2 [26], we applied a similar approach in the infection assay by pre-incubating SARS2-VLP with angiotensins for one hour at 37°C prior to infecting HEK293T-ACE2 target cells. As our preliminary data confirmed that SARS2-VLP infection is completed within 24 hours post-infection (data not shown), the VLP–angiotensin peptides mixture was removed at 24 hours post-infection, followed by a medium exchange after a washing step. To evaluate the dose-dependent effects of angiotensin peptides, we tested three concentrations: 40 nM, 80 nM, and 400LnM. Angiotensin II showed no effect on SARS-CoV-2 entry at any of the tested concentrations (Figure 3A). In contrast, angiotensin IV significantly increased viral infectivity at 40LnM, while higher concentrations (80 nM and 400 nM) led to a reduction in infectivity (Figure 3B), possibly due to cytotoxic or signaling effects. Notably, 40 nM angiotensin IV treatment consistently produced more than a twofold increase in luciferase activity across fifteen independent experiments, suggesting enhanced viral entry via the ACE2 receptor. However, the inhibitory effects observed at higher concentrations suggest that angiotensin IV may also exert antiviral effects that offset its entry-promoting activity. These findings highlight the unique dual role of angiotensin IV: enhancing viral entry at low concentrations predicted by increasing spike–ACE2 binding, while potentially initiating compensatory antiviral responses at higher concentrations. To further confirm that the angiotensin IV–mediated enhancement of SARS-CoV-2 target cell entry occurs specifically during the viral entry phase, rather than as a result of post-entry modulation of reporter gene expression, we treated HEK293T-ACE2 cells with 40LnM angiotensin IV at 24 hours post-infection, following viral removal and medium replacement (Figure 3C). This post-infection treatment did not significantly alter luciferase activity, supporting the conclusion that angiotensin IV primarily influences the viral entry stage.

**Figure 3.**
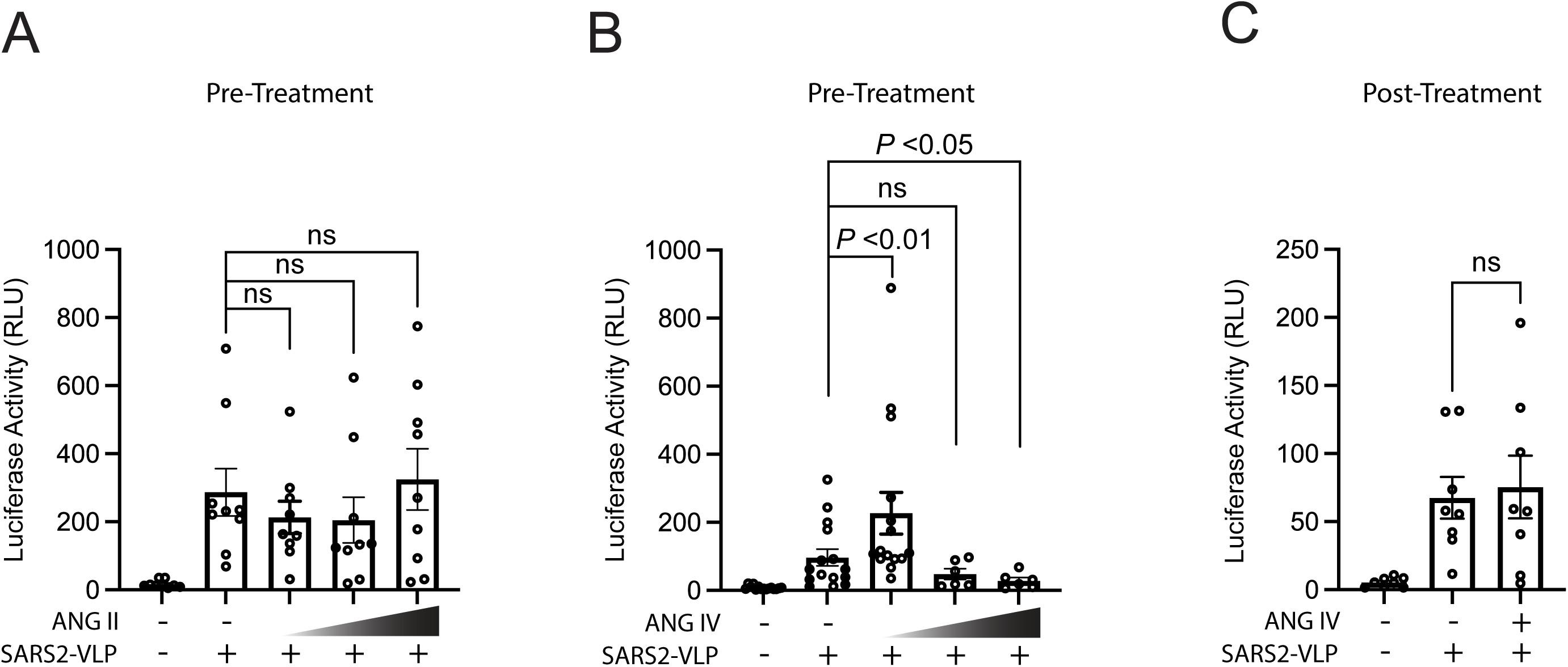
Effects of Angiotensin II and IV on the Viral Entry Step of SARS-CoV-2 Infection. SARS2-VLPs were pre-treated with angiotensin peptides at concentrations of 40LnM, 80LnM, and 400LnM prior to infection of HEK293T-ACE2 cells. **(A)** Angiotensin II treatment and **(B)** angiotensin IV treatment were assessed for their effects on viral entry. **(C)** To examine the post-entry effects, HEK293T-ACE2 cells were first infected with SARS2-VLPs and incubated for 24 hours. After removing residual virus by washing, cells were treated with 40LnM angiotensin IV for an additional 24 hours before measuring luciferase activity. Viral entry efficiency was quantified 48 hours post-infection using a luciferase reporter assay. Error bars represent the standard error of the mean. **(A)** For angiotensin II, 9 independent experiments were conducted at each concentration. **(B)** The 40LnM angiotensin IV group included 15 independent experiments, while the 80LnM and 400LnM groups included 6 independent experiments each. **(C)** A total of 8 independent experiments were performed for the post-entry 40 nM angiotensin IV treatment. Statistical significance was evaluated using the Wilcoxon matched-pairs signed-rank test: 9 matched pairs for angiotensin II compared to untreated samples and 15 matched pairs for the 40LnM angiotensin IV pretreatment group, 6 matched pairs for the 80LnM and 400LnM groups or 8 matched pairs for angiotensin IV post-treatment groups, respectively, compared to untreated samples.

### 3.2. Molecular Docking Analysis of Angiotensin Peptides and Spike Protein Interactions

Although angiotensin II is a well-known substrate of ACE2, prompting a natural interaction with the receptor, we sought to determine whether angiotensin IV also binds to ACE2. Structural simulations of angiotensin II and IV within the spike–ACE2 complex revealed that both peptides primarily associate with the enzymatic groove of ACE2 (Figure 4A [I] and [II]) and exhibit comparable binding affinities, suggesting that their affinity for ACE2 may exceed that of the spike protein (Table 1). However, angiotensin II did not enhance spike–ACE2 interaction [26], raising the possibility that if angiotensin IV binds to the same ACE2 site, it may similarly have no effect on spike–ACE2 binding. Based on this, we hypothesized that angiotensin IV may also interact directly with the spike protein, particularly its trimeric form, which is the native conformation present on intact virions. Interestingly, additional protein–ligand docking simulations showed that angiotensin IV preferentially binds to the S2 domain of the closed trimeric form of the spike protein (Figure 3B [I]). However, when simulated with the open-state conformation of the SARS-CoV-2 spike ectodomain, angiotensin IV bound in close proximity to the RBD of the S1 subunit, exhibiting increased binding affinity (Figure 3D [II] and Table 1).

**Figure 4:**
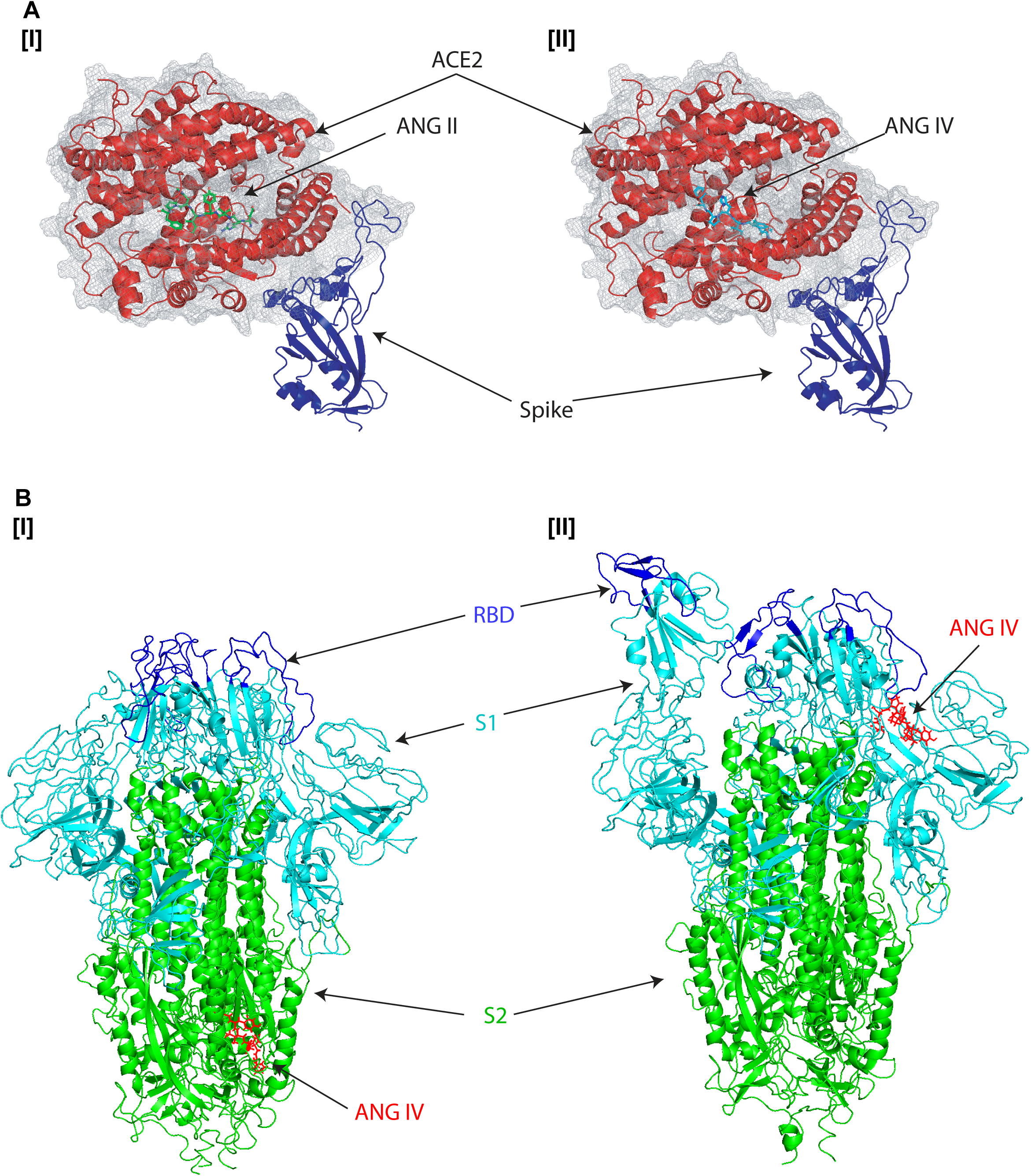
In silico docking models of angiotensin peptides with the SARS-CoV-2 spike protein and ACE2. **(A)** Predicted binding of **(I)** angiotensin II (green) and **(II)** angiotensin IV (cyan) to the SARS-CoV-2 spike receptor-binding domain (RBD, blue) in complex with ACE2 (red) based on the crystal structure (PDB: 6M0J), generated using DynamicBind. Both peptides were predicted to interact with the enzymatic groove of ACE2. Binding affinities are summarized in Table 1. **(B)** Simulated binding of angiotensin IV (red) to the SARS-CoV-2 spike trimer in its **(I)** closed (PDB: 6VXX) or **(II)** open (PDB: 6VYB) conformations. In these models, the RBD is shown in blue, the S1 domain in cyan, and the S2 domain in green.

**Table 1.**
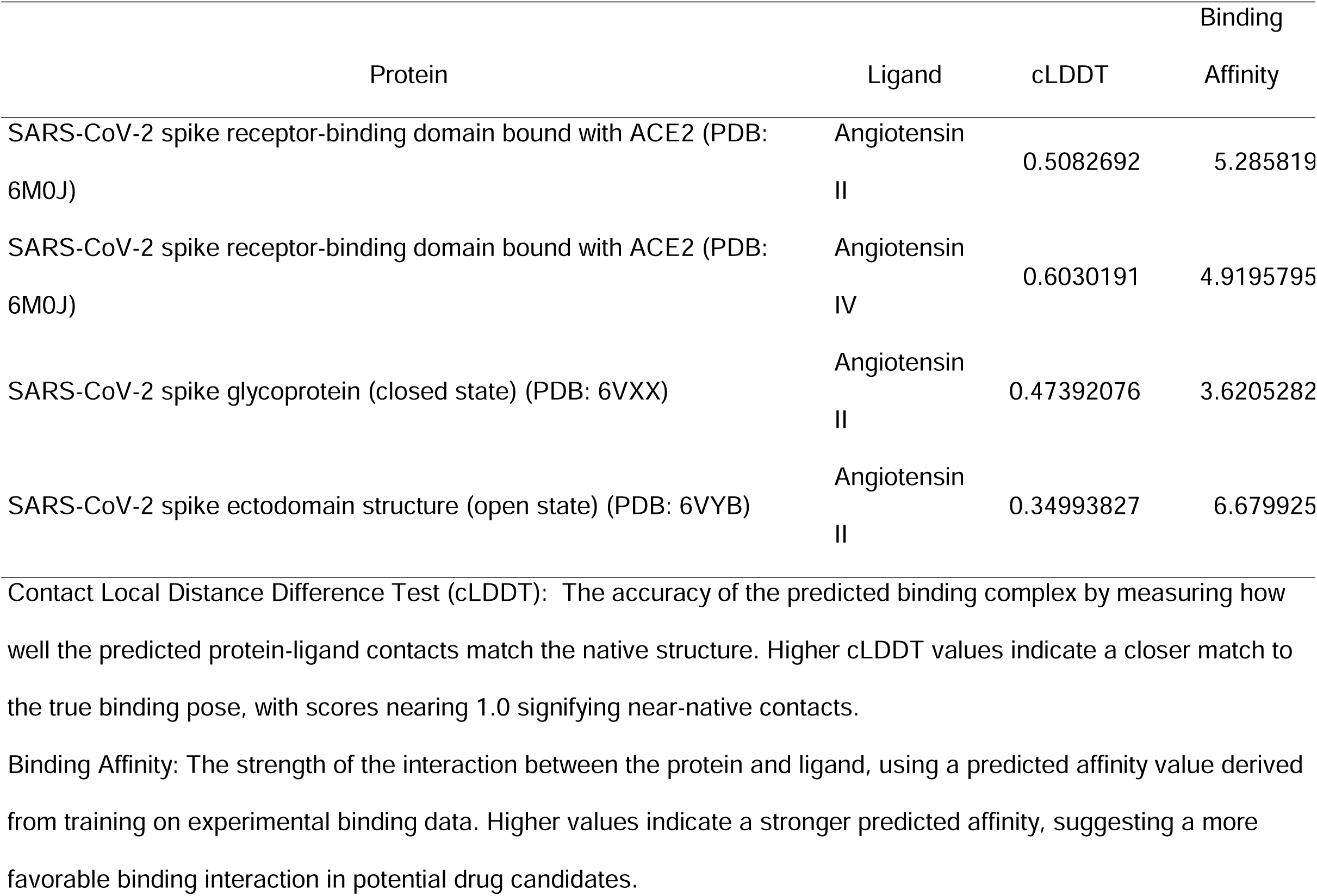
DynamicBind Metrics.

## 4. Discussion

This study demonstrates that specific downstream angiotensin peptides, particularly angiotensin IV, significantly influence SARS-CoV-2 entry into target cells via the ACE2 receptor (Figure 3B). While angiotensin II showed no measurable effect on spike–ACE2 binding, recent reports indicate that the shorter, metabolically derived peptides, angiotensin IV, significantly enhance binding affinity in ELISA-based assays [26]. This increased interaction may account for the transient rise in infectivity observed at lower concentrations of angiotensin IV in our single-round infection assays using VLPs (Figure 3B). Molecular docking simulations suggest that angiotensin IV interacts not only with the enzymatic groove of ACE2 but also binds to the S2 domain of the trimeric spike protein, distinct from the spike, ACE2 binding interface (Figure 4). Notably, angiotensin IV binds to the same domain of ACE2 as angiotensin II, which does not increase the virus infection. This suggests that the ability of angiotensin IV to increase binding may be attributed to its direct interaction with the spike protein. Since the predicted angiotensin IV binding site on the spike protein is located within the S2 domain, spatially distinct from the RBD in the S1 subunit, this separation suggests the potential for an allosteric mechanism. In this model, angiotensin IV may induce conformational changes in the spike protein that increase its affinity for ACE2, thereby facilitating viral entry. Furthermore, in silico docking simulations also indicate that angiotensin IV binds more tightly to the open conformation of the spike protein near the RBD than to ACE2 itself (Figure 4B and Table 1), supporting the notion that angiotensin IV preferentially interacts with the open state of the spike protein to enhance spike– ACE2 engagement. Importantly, these findings raise new questions about the physiological contexts in which angiotensin IV levels are altered, particularly regarding sex and age differences. Several studies have shown that RAS activity differs between males and females. Females tend to exhibit higher ACE2 expression in specific tissues compared to males, particularly in the renal tubulointerstitial compartment of the kidney [34]. This sex difference is thought to be partly driven by estrogen, which has been reported to upregulate ACE2 expression and may contribute to the observed disparities in ACE2 levels between sexes [18, 19]. This is also correlated with the fact that males tend to exhibit higher baseline levels of angiotensin II [20], while females may have relatively higher levels of angiotensin (1–7) and greater AT2R signaling, which are associated with vasoprotective and anti-inflammatory effects [21, 22]. Current research indicates that males and females have comparable susceptibility to SARS-CoV-2 infection [35], suggesting that sex-based differences in circulating angiotensin II levels may not significantly impact viral entry, which is consistent with our in vitro findings that angiotensin II does not affect SARS-CoV-2 cellular entry mechanisms (Figure 3A). However, males are at greater risk for severe COVID-19 outcomes, including higher rates of hospitalization, ICU admission, and mortality [36, 37], potentially due to elevated baseline levels of angiotensin II. While multiple biological factors likely contribute to these sex-based disparities, differences in angiotensin II concentrations may play a more prominent role in disease progression and severity rather than influencing initial viral infectivity.

While circulating levels of angiotensin II are well-documented, much less is known about angiotensin IV concentrations in circulation or within specific tissues, as it is a downstream metabolite generated through sequential enzymatic processing, including cleavage by APA. It is plausible that older adults and males, who generally experience reduced ACE2 expression and increased angiotensin II activity, may have altered conversion dynamics leading to elevated angiotensin IV levels in certain tissues, such as the lungs, heart, and kidneys, organs vulnerable to severe COVID-19. Angiotensin IV is likely present in both the CNS and peripheral tissues, as suggested by the widely distributed AT4R, however, its compartment-specific expression patterns and functional effects appear to differ between these regions [38–41]. In the CNS, angiotensin IV is generated locally and is abundantly expressed in regions such as the hippocampus, cortex, and hypothalamus, where it plays important roles in cognitive function, memory regulation, and neuroprotection [42, 43]. Those regions have been reported to exhibit structural and functional alterations in COVID-19 patients, including inflammation, neuronal injury, and disrupted neurogenesis, which may contribute to cognitive impairments in COVID-19, such as brain fog [44, 45]. In contrast, angiotensin IV is less abundant in plasma due to its rapid degradation and its role as a transient metabolic intermediate of angiotensin III, contributing only minimally to systemic vascular regulation [38]. Plasma angiotensin IV concentrations have been reported to vary widely, with one study indicating a median level of 39.71 ng/mL (81.6LnM) [27], while another reported a median of 8.6Lpg/mL (17.7 pM) [28], suggesting substantial variability across studies. Our in vitro data suggest that viral infectivity increases at 40LnM angiotensin IV, but higher concentrations beyond 80 nM reduce infectivity (Figure 3B). The median plasma concentration of angiotensin IV, based on limited two studies, is approximately 40LnM. While this estimate may lack precision, it suggests that under physiological conditions, angiotensin IV is more likely to enhance SARS-CoV-2 infection rather than inhibit it.

Studies have demonstrated that SARS-CoV-2 can directly infect human microglial cells, leading to their activation and triggering a proinflammatory response, including the release of IL-1β, IL-6, and TNF-α, which are associated with neuroinflammation and may underlie the neurological symptoms observed in COVID-19 [46], which closely resembles the pathogenic mechanisms seen in HIV associated neurocognitive disorders [47]. Given our findings that angiotensin IV facilitates viral entry, individuals with inherently higher local or systemic levels of angiotensin IV, particularly within the CNS, may be more susceptible to enhanced SARS-CoV-2 entry into neurons or microglial cells. This increased susceptibility could contribute to more severe neuroinvasion and may underline the neurocognitive impairments, such as brain fog, commonly observed in COVID-19 patients. A large cohort study reported that females are at significantly higher risk of developing neurological symptoms associated with COVID-19, such as fatigue, headache, brain fog, depression, and anosmia, compared to males [48]. The risk ratios for these symptoms in females range from 1.37 to 1.61, indicating a 37% to 61% higher risk than males for specific symptoms [49]. Although there is currently no direct evidence of sex-based differences in angiotensin IV levels within the CNS, such differences may contribute to the observed sex disparities in COVID-19-related neurological complications. Further clinical research is needed to quantify angiotensin peptide concentrations in both plasma and the CNS to clarify their potential role in sex-specific disease manifestations.

Although angiotensin IV does not universally reduce cell viability in vitro, it has been reported to decrease proliferation and viability in certain tumor-derived cell lines, such as GH3 cells, at concentrations as low as 10 nM and 0.1 nM following 72 hours of treatment, potentially through mechanisms independent of classical angiotensin receptors [50]. While these findings suggest that the reduced viral infectivity observed at higher concentrations of angiotensin IV may be partially due to cytotoxic effects in HEK293T-ACE2 cells, it is noteworthy that angiotensin II, despite reducing GH3 cell viability at 0.1 nM, did not impact viral entry even at 400LnM (Figure 3A). This implies that HEK293T-ACE2 cells may be relatively resistant to angiotensin-induced cytotoxicity at the concentrations tested. Therefore, the reduction in infectivity observed with high concentrations of angiotensin IV may be more likely attributed to a modest antiviral response rather than direct cytotoxicity. Given the wide variability in reported plasma concentrations of angiotensin IV, physiological levels may be insufficient to exert a direct antiviral effect. However, future studies identifying antiviral mechanisms triggered by angiotensin IV could pave the way for the development of novel anti-SARS-CoV-2 therapeutics. In the current context, the primary role of angiotensin IV may lie in enhancing viral infectivity by increasing spike–ACE2 binding affinity.

In summary, our study identifies angiotensin IV as a potential endogenous enhancer of SARS-CoV-2 cell entry via the ACE2 receptor. These findings open new avenues for understanding how RAS imbalance, particularly involving downstream metabolites, may contribute to COVID-19 susceptibility and pathogenesis. Future research should explore angiotensin IV expression profiles across sexes, ages, and disease states, as well as the therapeutic potential of targeting angiotensin IV–related pathways to mitigate viral infectivity.

## Author Contributions

S.G. and T.H. contributed equally to this work and should be considered co-first authors. Y.J.S. and T.I. conceptualized the study, and T.I. designed it. S.G., T.H., H.T., D.M., G.H., A.D., and T.I. carried out the experiments. Data analysis was conducted by S.G., T.H., H.T., D.M., G.H., A.D., J.C., C.I., and T.I. The manuscript was written by S.G., T.H., and T.I. and revised by all authors. Y.J.S. and T.I. provided financial support for the research. All authors have read and agreed to the submission version of the manuscript.

## Institutional Review Board Statement

This study has been approved by the American University Institutional Biosafety Committee since August 08, 2023.

## Acknowledgments

We greatly appreciate the contributions of the American University undergraduate student, Cecelia Cropp, for her active suggestions and comments on this manuscript. We express our gratitude to the administrators and leadership of the authors’ institutions, Georgetown University and American University, for their continued support of this manuscript submission.

## References

1. Zhang, J. J.; Dong, X.; Liu, G. H.; Gao, Y. D., Risk and Protective Factors for COVID-19 Morbidity, Severity, and Mortality. Clin Rev Allergy Immunol 2023, 64, (1), 90–107.

2. Chakravarty, D.; Das Sarma, J., Murine-beta-coronavirus-induced neuropathogenesis sheds light on CNS pathobiology of SARS-CoV2. J Neurovirol 2021, 27, (2), 197–216.

3. Zhang, P.; Yu, L.; Dong, J.; Liu, Y.; Zhang, L.; Liang, P.; Wang, L.; Chen, B.; Huang, L.; Song, C., Cellular poly(C) binding protein 2 interacts with porcine epidemic diarrhea virus papain-like protease 1 and supports viral replication. Vet Microbiol 2020, 247, 108793.

4. Ashraf, U. M.; Abokor, A. A.; Edwards, J. M.; Waigi, E. W.; Royfman, R. S.; Hasan, S. A.; Smedlund, K. B.; Hardy, A. M. G.; Chakravarti, R.; Koch, L. G., SARS-CoV-2, ACE2 expression, and systemic organ invasion. Physiol Genomics 2021, 53, (2), 51–60.

5. Antony, P.; Vijayan, R., Role of SARS-CoV-2 and ACE2 variations in COVID-19. Biomed J 2021, 44, (3), 235–244.

6. Yang, J.; Chen, T.; Zhou, Y., Mediators of SARS-CoV-2 entry are preferentially enriched in cardiomyocytes. Hereditas 2021, 158, (1), 4.

7. Rotondi, M.; Coperchini, F.; Ricci, G.; Denegri, M.; Croce, L.; Ngnitejeu, S. T.; Villani, L.; Magri, F.; Latrofa, F.; Chiovato, L., Detection of SARS-COV-2 receptor ACE-2 mRNA in thyroid cells: a clue for COVID-19-related subacute thyroiditis. J Endocrinol Invest 2021, 44, (5), 1085–1090.

8. Bian, J.; Li, Z., Angiotensin-converting enzyme 2 (ACE2): SARS-CoV-2 receptor and RAS modulator. Acta Pharm Sin B 2021, 11, (1), 1–12.

9. Santos, R. A. S.; Sampaio, W. O.; Alzamora, A. C.; Motta-Santos, D.; Alenina, N.; Bader, M.; Campagnole-Santos, M. J., The ACE2/Angiotensin-(1-7)/MAS Axis of the Renin-Angiotensin System: Focus on Angiotensin-(1-7). Physiol Rev 2018, 98, (1), 505–553.

10. Delaitre, C.; Boisbrun, M.; Lecat, S.; Dupuis, F., Targeting the Angiotensin II Type 1 Receptor in Cerebrovascular Diseases: Biased Signaling Raises New Hopes. Int J Mol Sci 2021, 22, (13).

11. Wysocki, J.; Ye, M.; Batlle, D., Plasma and Kidney Angiotensin Peptides: Importance of the Aminopeptidase A/Angiotensin III Axis. Am J Hypertens 2015, 28, (12), 1418–26.

12. Kemp, B. A.; Bell, J. F.; Rottkamp, D. M.; Howell, N. L.; Shao, W.; Navar, L. G.; Padia, S. H.; Carey, R. M., Intrarenal angiotensin III is the predominant agonist for proximal tubule angiotensin type 2 receptors. Hypertension 2012, 60, (2), 387–95.

13. Padia, S. H.; Kemp, B. A.; Howell, N. L.; Fournie-Zaluski, M. C.; Roques, B. P.; Carey, R. M., Conversion of renal angiotensin II to angiotensin III is critical for AT2 receptor-mediated natriuresis in rats. Hypertension 2008, 51, (2), 460–5.

14. Park, B. M.; Cha, S. A.; Lee, S. H.; Kim, S. H., Angiotensin IV protects cardiac reperfusion injury by inhibiting apoptosis and inflammation via AT4R in rats. Peptides 2016, 79, 66–74.

15. Zulli, A.; Burrell, L. M.; Buxton, B. F.; Hare, D. L., ACE2 and AT4R are present in diseased human blood vessels. Eur J Histochem 2008, 52, (1), 39–44.

16. Gul, R.; Kim, U. H.; Alfadda, A. A., Renin-angiotensin system at the interface of COVID-19 infection. Eur J Pharmacol 2021, 890, 173656.

17. Wigen, J.; Lofdahl, A.; Bjermer, L.; Elowsson-Rendin, L.; Westergren-Thorsson, G., Converging pathways in pulmonary fibrosis and Covid-19 - The fibrotic link to disease severity. Respir Med X 2020, 2, 100023.

18. Bukowska, A.; Spiller, L.; Wolke, C.; Lendeckel, U.; Weinert, S.; Hoffmann, J.; Bornfleth, P.; Kutschka, I.; Gardemann, A.; Isermann, B.; Goette, A., Protective regulation of the ACE2/ACE gene expression by estrogen in human atrial tissue from elderly men. Exp Biol Med (Maywood) 2017, 242, (14), 1412–1423.

19. Youn, J. Y.; Zhang, Y.; Wu, Y.; Cannesson, M.; Cai, H., Therapeutic application of estrogen for COVID-19: Attenuation of SARS-CoV-2 spike protein and IL-6 stimulated, ACE2-dependent NOX2 activation, ROS production and MCP-1 upregulation in endothelial cells. Redox Biol 2021, 46, 102099.

20. Liu, Y.; Cui, Y.; Zhou, Z.; Liu, B.; Liu, Z.; Li, G., Relationship between Angiotensin II, Vascular Endothelial Growth Factor, and Arteriosclerosis Obliterans. Dis Markers 2023, 2023, 1316821.

21. Vinh, A.; Widdop, R. E.; Chai, S. Y.; Gaspari, T. A., Angiotensin IV-evoked vasoprotection is conserved in advanced atheroma. Atherosclerosis 2008, 200, (1), 37–44.

22. Bhat, S. A.; Fatima, Z.; Sood, A.; Shukla, R.; Hanif, K., The Protective Effects of AT2R Agonist, CGP42112A, Against Angiotensin II-Induced Oxidative Stress and Inflammatory Response in Astrocytes: Role of AT2R/PP2A/NFkappaB/ROS Signaling. Neurotox Res 2021, 39, (6), 1991–2006.

23. Gomez, J. M. D.; Du-Fay-de-Lavallaz, J. M.; Fugar, S.; Sarau, A.; Simmons, J. A.; Clark, B.; Sanghani, R. M.; Aggarwal, N. T.; Williams, K. A.; Doukky, R.; Volgman, A. S., Sex Differences in COVID-19 Hospitalization and Mortality. J Womens Health (Larchmt) 2021, 30, (5), 646–653.

24. Nguyen, N. T.; Chinn, J.; De Ferrante, M.; Kirby, K. A.; Hohmann, S. F.; Amin, A., Male gender is a predictor of higher mortality in hospitalized adults with COVID-19. PLoS One 2021, 16, (7), e0254066.

25. Wilcox, T.; Smilowitz, N. R.; Seda, B.; Xia, Y.; Hochman, J.; Berger, J. S., Sex Differences in Thrombosis and Mortality in Patients Hospitalized for COVID-19. Am J Cardiol 2022, 170, 112–117.

26. Oliveira, K. X.; Suzuki, Y. J., Angiotensin peptides enhance SARS-CoV-2 spike protein binding to its host cell receptors. bioRxiv 2024.

27. Martyniak, A.; Drozdz, D.; Tomasik, P. J., Classical and Alternative Pathways of the Renin-Angiotensin-Aldosterone System in Regulating Blood Pressure in Hypertension and Obese Adolescents. Biomedicines 2024, 12, (3).

28. Shibasaki, Y.; Mori, Y.; Tsutumi, Y.; Masaki, H.; Sakamoto, K.; Murasawa, S.; Maruyama, K.; Moriguchi, Y.; Tanaka, Y.; Iwasaka, T.; Inada, M.; Matsubara, H., Differential kinetics of circulating angiotensin IV and II after treatment with angiotensin II type 1 receptor antagonist and their plasma levels in patients with chronic renal failure. Clin Nephrol 1999, 51, (2), 83–91.

29. Syed, A. M.; Taha, T. Y.; Tabata, T.; Chen, I. P.; Ciling, A.; Khalid, M. M.; Sreekumar, B.; Chen, P. Y.; Hayashi, J. M.; Soczek, K. M.; Ott, M.; Doudna, J. A., Rapid assessment of SARS-CoV-2-evolved variants using virus-like particles. Science 2021, 374, (6575), 1626–1632.

30. Shang, J.; Ye, G.; Shi, K.; Wan, Y.; Luo, C.; Aihara, H.; Geng, Q.; Auerbach, A.; Li, F., Structural basis of receptor recognition by SARS-CoV-2. Nature 2020, 581, (7807), 221–224.

31. Lan, J.; Ge, J.; Yu, J.; Shan, S.; Zhou, H.; Fan, S.; Zhang, Q.; Shi, X.; Wang, Q.; Zhang, L.; Wang, X., Structure of the SARS-CoV-2 spike receptor-binding domain bound to the ACE2 receptor. Nature 2020, 581, (7807), 215–220.

32. Walls, A. C.; Park, Y. J.; Tortorici, M. A.; Wall, A.; McGuire, A. T.; Veesler, D., Structure, Function, and Antigenicity of the SARS-CoV-2 Spike Glycoprotein. Cell 2020, 181, (2), 281–292 e6.

33. Lu, W.; Zhang, J.; Huang, W.; Zhang, Z.; Jia, X.; Wang, Z.; Shi, L.; Li, C.; Wolynes, P. G.; Zheng, S., DynamicBind: predicting ligand-specific protein-ligand complex structure with a deep equivariant generative model. Nat Commun 2024, 15, (1), 1071.

34. Maksimowski, N. A.; Scholey, J. W.; Williams, V. R.; Nephrotic Syndrome Study, N., Sex and kidney ACE2 expression in primary focal segmental glomerulosclerosis: A NEPTUNE study. PLoS One 2021, 16, (6), e0252758.

35. Scully, E. P.; Schumock, G.; Fu, M.; Massaccesi, G.; Muschelli, J.; Betz, J.; Klein, E. Y.; West, N. E.; Robinson, M.; Garibaldi, B. T.; Bandeen-Roche, K.; Zeger, S.; Klein, S. L.; Gupta, A., Sex and Gender Differences in Testing, Hospital Admission, Clinical Presentation, and Drivers of Severe Outcomes From COVID-19. Open Forum Infect Dis 2021, 8, (9), ofab448.

36. Peckham, H.; de Gruijter, N. M.; Raine, C.; Radziszewska, A.; Ciurtin, C.; Wedderburn, L. R.; Rosser, E. C.; Webb, K.; Deakin, C. T., Male sex identified by global COVID-19 meta-analysis as a risk factor for death and ITU admission. Nat Commun 2020, 11, (1), 6317.

37. Sieurin, J.; Branden, G.; Magnusson, C.; Hergens, M. P.; Kosidou, K., A population-based cohort study of sex and risk of severe outcomes in covid-19. Eur J Epidemiol 2022, 37, (11), 1159–1169.

38. Wright, J. W.; Krebs, L. T.; Stobb, J. W.; Harding, J. W., The angiotensin IV system: functional implications. Front Neuroendocrinol 1995, 16, (1), 23–52.

39. Zhuo, J.; Moeller, I.; Jenkins, T.; Chai, S. Y.; Allen, A. M.; Ohishi, M.; Mendelsohn, F. A., Mapping tissue angiotensin-converting enzyme and angiotensin AT1, AT2 and AT4 receptors. J Hypertens 1998, 16, (12 Pt 2), 2027–37.

40. Chai, S. Y.; Bastias, M. A.; Clune, E. F.; Matsacos, D. J.; Mustafa, T.; Lee, J. H.; McDowall, S. G.; Paxinos, G.; Mendelsohn, F. A.; Albiston, A. L., Distribution of angiotensin IV binding sites (AT4 receptor) in the human forebrain, midbrain and pons as visualised by in vitro receptor autoradiography. J Chem Neuroanat 2000, 20, (3-4), 339–48.

41. de Gasparo, M.; Catt, K. J.; Inagami, T.; Wright, J. W.; Unger, T., International union of pharmacology. XXIII. The angiotensin II receptors. Pharmacol Rev 2000, 52, (3), 415–72.

42. Gard, P. R., Cognitive-enhancing effects of angiotensin IV. BMC Neurosci 2008, 9 Suppl 2, (Suppl 2), S15.

43. Wright, J. W.; Harding, J. W., Contributions by the Brain Renin-Angiotensin System to Memory, Cognition, and Alzheimer’s Disease. J Alzheimers Dis 2019, 67, (2), 469–480.

44. Lu, Y.; Li, X.; Geng, D.; Mei, N.; Wu, P. Y.; Huang, C. C.; Jia, T.; Zhao, Y.; Wang, D.; Xiao, A.; Yin, B., Cerebral Micro-Structural Changes in COVID-19 Patients - An MRI-based 3-month Follow-up Study. EClinicalMedicine 2020, 25, 100484.

45. Zhou, S.; Wei, T.; Liu, X.; Liu, Y.; Song, W.; Que, X.; Xing, Y.; Wang, Z.; Tang, Y., Causal effects of COVID-19 on structural changes in specific brain regions: a Mendelian randomization study. BMC Med 2023, 21, (1), 261.

46. Jeong, G. U.; Lyu, J.; Kim, K. D.; Chung, Y. C.; Yoon, G. Y.; Lee, S.; Hwang, I.; Shin, W. H.; Ko, J.; Lee, J. Y.; Kwon, Y. C., SARS-CoV-2 Infection of Microglia Elicits Proinflammatory Activation and Apoptotic Cell Death. Microbiol Spectr 2022, 10, (3), e0109122.

47. Gagliardi, S.; Hotchkin, T.; Hillmer, G.; Engelbride, M.; Diggs, A.; Tibebe, H.; Izumi, C.; Sullivan, C.; Cropp, C.; Lantz, O., Oxidative Stress in HIV-Associated Neurodegeneration: Mechanisms of Pathogenesis and Therapeutic Targets. 2025.

48. Gorenshtein, A.; Leibovitch, L.; Liba, T.; Stern, S.; Stern, Y., Gender Disparities in Neurological Symptoms of Long COVID: A Systematic Review and Meta-Analysis. Neuroepidemiology 2024, 1–15.

49. Bai, F.; Tomasoni, D.; Falcinella, C.; Barbanotti, D.; Castoldi, R.; Mule, G.; Augello, M.; Mondatore, D.; Allegrini, M.; Cona, A.; Tesoro, D.; Tagliaferri, G.; Vigano, O.; Suardi, E.; Tincati, C.; Beringheli, T.; Varisco, B.; Battistini, C. L.; Piscopo, K.; Vegni, E.; Tavelli, A.; Terzoni, S.; Marchetti, G.; Monforte, A. D., Female gender is associated with long COVID syndrome: a prospective cohort study. Clin Microbiol Infect 2022, 28, (4), 611 e9–611 e16.

50. Ptasinska-Wnuk, D.; Lawnicka, H.; Mucha, S.; Kunert-Radek, J.; Pawlikowski, M.; Stepien, H., Angiotensins inhibit cell growth in GH3 lactosomatotroph pituitary tumor cell culture: a possible involvement of the p44/42 and p38 MAPK pathways. ScientificWorldJournal 2012, 2012, 189290.

